# The ATLAS Penalty: Auxiliary-Transformed Location-Aware Smoothing with Applications to Spatial Transcriptomics

**DOI:** 10.64898/2026.05.18.725545

**Authors:** Qian Tang, Eric C. Chi, Wenyi Wang

## Abstract

We address the problem of fitting a collection of location-specific models under a spatial smoothness assumption. Existing approaches penalize roughness in the model parameters directly, an assumption that breaks down when smoothness is a function of parameters and auxiliary covariates rather than the parameters themselves. Our framework, the Auxiliary-Transformed Location-Aware Smoothing (ATLAS) penalty, generalizes spatial smoothness by penalizing roughness in transformations of model parameters using auxiliary information. As a concrete case study, we develop a spatially smooth deconvolution model for spatial transcriptomics that estimates tumor mixing coefficients from thousands of spots distributed on a single tissue slide. To handle the computational challenges posed by the nonlinear likelihood, nonsmooth nonconvex penalty, and spatially coupled estimation, we propose an alternating direction method of multipliers (ADMM) algorithm. Through simulation studies, we demonstrate that our framework provides substantially better spatial domain detection than approaches that smooth model parameters directly, with particularly strong gains when auxiliary covariates carry calibrated spatial structure.

## 1 Introduction

We study the problem of fitting a collection of location-specific models under spatial smoothness. Formally, we observe response vectors *Y*_*i*_ ∈R^*p*^ that follow a law *F* (·| ***θ***_*i*_) for *i* ∈ [*S*] = {1, …, *S*} spots or locations. The 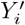 *s* are independent conditional on ***θ***_*i*_. The goal is to recover location model’s ***θ***_*i*_ based on the observed *Y*_*i*_ and a spatial smoothness assumption on the ***θ***_*i*_. This task is central to a wide and diverse spectrum of domains including but not limited to neuroimaging [Hu and Allen, 2015], disease mapping [MacNab, 2011, Wang and Rodríguez, 2014], and air quality monitoring [Sun et al., 2016]. In all these problems, we wish to fit models at each location that respects spatial smoothness, as responses at adjacent locations are typically correlated.

Existing approaches, e.g., local-aggregate models Hu and Allen [2015] and network lasso Hallac et al. [2015], address this by penalizing roughness in the model parameters directly. The corresponding penalized maximum likelihood estimation problem is

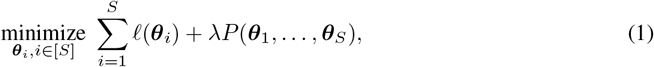

where *𝓁*(***θ***_*i*_) is a negative log-likelihood at the *i*th location and *P* is a regularizer that penalizes variation or roughness in the ***θ***_*i*_, and *λ* is a nonnegative tuning parameter that trades off model fit quantified by the likelihood and model complexity quantified by the penalty.

Problem (1) penalizes roughness in ***θ***_*i*_ directly. In some cases, however, this assumption breaks down. For example, in spatial transcriptomics, the difference between tumor transcript proportion and tumor cell purity is a biologically meaningful signal that clusters spatially near tumor-stromal boundaries [Montierth et al., 2025] – a signal that smoothing location model parameters independently cannot fully capture (**Appendix**). Consequently, we propose a generalized spatial smoothness framework that moves beyond directly penalizing parameter roughness by penalizing roughness on *h*_*i*_(***θ***_*i*_) = *h*(***θ***_*i*_; ***ψ***_*i*_) — a transformation shared across locations but parameterized by location-specific auxiliary variables ***ψ***_*i*_. Specifically, we propose an alternative penalized maximum likelihood estimation problem

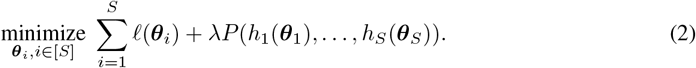

We call this the Auxiliary-Transformed Location-Aware Smoothing (ATLAS) penalty. The availability of additional location dependent parameters ***ψ***_*i*_ is quite common. These are often measured quantities that do not play an explicit role in the likelihood. Biologically, ATLAS penalty is designed to impose spatial smoothing over the desired biological signals directly, when clustering over ***θ***_**i**_^′^*s* or ***ψ***^′^_*i*_*s* separately is more susceptible to technical artifacts, e.g., clustering separately is not sufficient to classify spots as specific tumor microenvironment niches [Montierth et al., 2025]. While the proposed framework is general, as a concrete case study, we develop a spatially smooth deconvolution model for spatial transcriptomics.

## 2 ATLAS for Spot-level Deconvolution in Spatial Transcriptomics

Cancer is a complex disease at the cellular level. Tumors exhibit a high degree of variability arising from genetic, phenotypic, and microenvironmental heterogeneity within the local tumor microenvironment, impacting progression, spread and therapeutic resistance [Marusyk et al., 2012, 2020]. Moreover, malignant cells exhibit phenotypic plasticity, transitioning between cell states, further contributing to intratumor diversity [Meacham and Morrison, 2013, Bhat et al., 2024]. Indeed, tumor cell transcriptional output varies substantially across tumors and tumor cell states [LaFave et al., 2020, Cao et al., 2022]. At the single-cell level, transcriptional diversity within a tumor correlates with developmental plasticity and stemness [Gulati et al., 2020], further amplifying this variability. Consequently, tumor-cell specific transcript proportion in tissues can serve as a potential biomarker for disease progression prediction and refined tumor stratification, with implications for more targeted treatment strategies [Cao et al., 2022].

We consider a spatial transcriptomic (ST) sample with *S* spots and *G* genes. Let *Y*_*ig*_ denote the observed expression count for gene *g* at spot *i*, for *i* = 1, …, *S* and *g* = 1, …, *G*. We model each observed count as the sum of two latent components corresponding to tumor and non-tumor contributions:

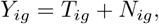

where *T*_*ig*_ denotes the tumor-derived count and *N*_*ig*_ denotes the non-tumor-derived count. Conditional on the model parameters, the two latent components are assumed to be independent.

For each spot *i*, let *π*_*i*_ ∈ (0, 1) denote the spot-specific tumor mixing coefficient. For each gene *g*, let *µ*_*tg*_ and *µ*_*ng*_ denote the baseline expression levels of the tumor and non-tumor components, respectively. We assume that the latent tumor and non-tumor counts follow negative binomial distributions,

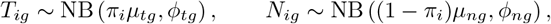

where *ϕ*_*tg*_ and *ϕ*_*ng*_ are gene-specific inverse-dispersion parameters for the tumor and non-tumor components, respectively.

Since *T*_*ig*_ and *N*_*ig*_ are latent and only their sum *Y*_*ig*_ = *T*_*ig*_ +*N*_*ig*_ is observed, the marginal distribution of *Y*_*ig*_ is the convolution of the two negative binomial components:

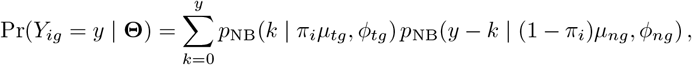

where **Θ** = (***π, µ***_*t*_, ***µ***_*n*_, ***ϕ***_*t*_, ***ϕ***_*n*_) denotes the complete set of parameters. Here *p*_NB_(·| *m, ϕ*) denotes the negative binomial probability mass function with mean *m* and inverse-dispersion parameter *ϕ*:

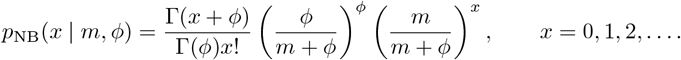

Assuming conditional independence across spots and genes, the observed-data negative log-likelihood is

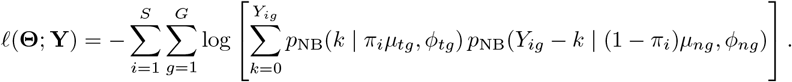

Montierth et al. [2025] used experimentally mixed benchmark data to validate the accuracy of 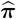 using maximum likelihood estimation, without accounting for any spatial relationship. The resulting 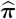 however, showed the best spatial clustering of spots when adjusted further by the Hematoxylin and Eosin-stained tissue image-based tumor cell proportions ***ρ***_*i*_, hence motivating spatial smoothing using *h*_*i*_(*π*_*i*_) = *π*_*i*_− *ρ*_*i*_ (**Appendix: Figure S1**). Given the gene-specific tumor and non-tumor parameters (***µ***_*t*_, ***µ***_*n*_, ***ϕ***_*t*_, ***ϕ***_*n*_), we estimate the spot-specific tumor mixing coefficients by minimizing a spatially penalized negative observed-data log-likelihood:

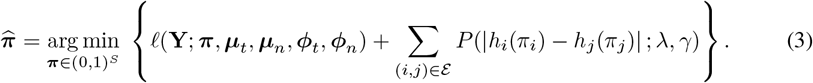

Here ε denotes the set of neighboring spot pairs induced by the spatial adjacency graph, with ε⊆{ (*i, j*) : 1≤ *i < j* ≤*S*}. The penalty is imposed on the transformed spatial signal *h*_*i*_(*π*_*i*_), rather than directly on the raw mixing coefficient *π*_*i*_. In this work, we consider the affine transformation

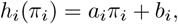

where *a*_*i*_ and *b*_*i*_ are prespecified location-specific constants. These constants allow auxiliary spatial information, such as calibrated tumor transcript proportions or histology-derived scores, to determine the scale on which spatial smoothness is imposed. In the absence of auxiliary information, one may set *a*_*i*_ = 1 and *b*_*i*_ = 0, in which case *h*_*i*_(*π*_*i*_) = *π*_*i*_ and the penalty reduces to direct spatial smoothing of the tumor mixing proportions.

For the penalty function *P*, we adopt the minimax concave penalty (MCP) [Zhang, 2010]:

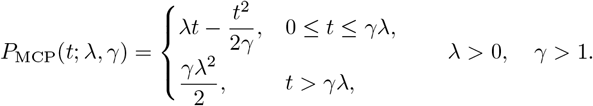

Here, *λ* controls the strength of spatial smoothing, while *γ* controls the concavity of the penalty. The MCP penalty shrinks small neighboring differences toward zero while leaving large differences unchanged. In the context of our problem, it promotes spatial homogeneity within domains and preserves sharp boundaries between distinct tissue regions.

### Remark

We justify using an affine *h*_*i*_(*π*_*i*_) = *a*_*i*_*π*_*i*_ + *b*_*i*_ as a first-order Taylor approximation of a smooth transformation around a spot-specific reference. As we will see in our experiments, although relatively simple, the affine model can effectively incorporate auxiliary information to improve segmentation. Moreover, the affine model also expedites computation with our proposed ADMM algorithm. We address more general *h*_*i*_ in the discussion.

## 3 Estimation Algorithm

Direct optimization of the objective function formulated in (3) poses significant computational challenges due to three primary factors. First, the observed-data likelihood exhibits a non-linear dependence on ***π***, arising from the convolution of the tumor and non-tumor negative binomial components. Second, the Minimax Concave Penalty (MCP) is inherently non-smooth and non-convex. Finally, the graph-based regularization induces structural coupling among the transformed mixing proportions across adjacent spatial locations. These characteristics collectively preclude direct, separable optimization over the spatial domain. To circumvent these intractable features, we employ a variable splitting strategy and optimize the resulting constrained formulation via the Alternating Direction Method of Multipliers (ADMM) [Boyd et al., 2011].

For ease of notation, we define the negative log-likelihood as *f* (***π***) = *𝓁* (**Y**; ***π, µ***_*t*_, ***µ***_*n*_, ***ϕ***_*t*_, ***ϕ***_*n*_). Let **A** = diag(*a*_1_, …, *a*_*S*_) and **b** = (*b*_1_, …, *b*_*S*_)^T^. The transformed spatial signal is formulated as **x** = **A*π*** + **b**, with the *i*-th component given by *x*_*i*_ = *h*_*i*_(*π*_*i*_) = *a*_*i*_*π*_*i*_ + *b*_*i*_. Furthermore, let **D** ∈ **ℝ**^*M* ×*S*^, where *M* =|ε|, denote the oriented edge-incidence matrix associated with the spatial graph. For an arbitrary edge *e* = (*i, j*) ∈ ε, the corresponding row in **D** is constructed such that (**Dx**)_*e*_ = *x*_*i*_ − *x*_*j*_. The orientation of the edges can be arbitrarily assigned, as the penalty function depends strictly on the absolute edgewise differences. By introducing the auxiliary variable **z** = **Dx**, the spatial penalty can be expressed as 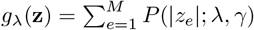. Consequently, problem (3) can be equivalently recast as the following constrained optimization problem:

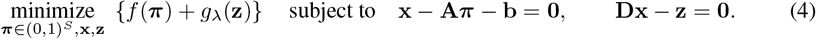

Introducing the penalty parameters *σ*_1_, *σ*_2_ *>* 0, the scaled augmented Lagrangian associated with (4) is formulated as:

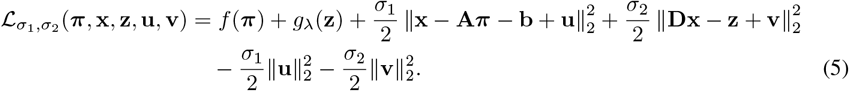

Here **u** ∈ℝ^*S*^ and **v** ∈ℝ^*M*^ are the Lagrangian multipliers and ∥· ∥_2_ denotes the Euclidean norm. ADMM minimizes the augmented Lagrangian one block of variables at a time. This yields the algorithm

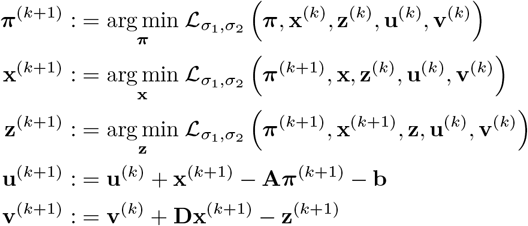

Minimization with respect to ***π*** entails solving the following subproblem:

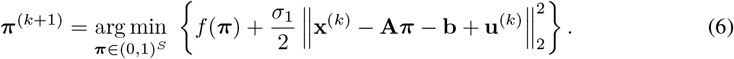

Since the observed-data likelihood factorizes over spots and the quadratic augmented Lagrangian term is separable in *π*_*i*_, the problem in (6) decomposes into *S* independent one-dimensional optimization problems. Specifically, for each spot *i*, we solve

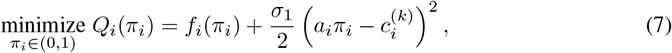

Where 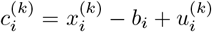. To enforce the constraint *π*_*i*_ ∈ (0, 1), we reparameterize the scalar problem using the logit transformation 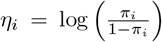 . and. 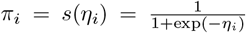 Each one-dimensional subproblem is then optimized with respect to the unconstrained variable *η*_*i*_ using Newton’s method with Armijo backtracking line search.

The update for **x** is obtained by minimizing the unconstrained quadratic program:

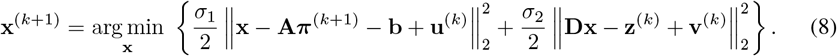

The first-order optimality condition yields the sparse linear system

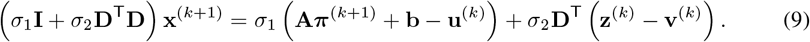

The coefficient matrix, *σ*_1_**I** + *σ*_2_**D**^T^**D**, is a shifted graph Laplacian that is strictly symmetric positive definite for any *σ*_1_ *>* 0, thereby ensuring a unique solution to the linear system. A key computational advantage is that this matrix is invariant to both the ADMM iteration index *k* and the regularization parameter *λ*. Consequently, its sparse Cholesky factorization can be pre-computed once and efficiently reused throughout all ADMM iterations and across the entire regularization path.

For the auxiliary edge-difference variable **z**, the ADMM update is

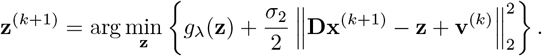

Define 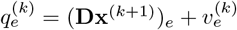. Then, for each edge *e* = 1, …, *M*, the update is

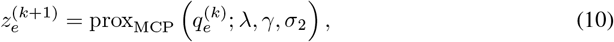

where the MCP proximal operator is defined by

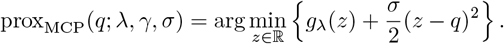

### Lemma 3.1

*Suppose λ >* 0, *γ >* 1, *and σ >* 1*/γ. Then, for any q* ∈ℝ, *the scalar optimization problem*(10) *has the closed-form solution*

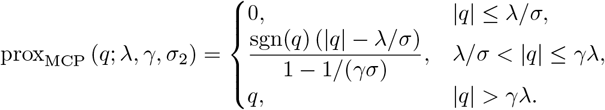

Additional details and the complete algorithm are provided in the **Appendix**.

## 4 Simulation Studies

We compare the proposed ATLAS estimator, which we denote by *h*(*π*)-MCP, with three alternative strategies for spatial domain detection, i.e. clustering accuracy of groups of spots/locations. The first baseline is direct spatial smoothing of the tumor mixing proportion, which assumes the spatial arrangement of spots is strongly associated with the spot-level tumor mixing proportions, denoted by *π*-MCP. This method corresponds to setting *a*_*i*_ = 1 and *b*_*i*_ = 0, so that *h*_*i*_(*π*_*i*_) = *π*_*i*_. The second comparison method uses only the auxiliary calibration quantity for domain detection. Specifically, spatial domains are inferred by applying k-means clustering to the auxiliary quantity. We refer to this method as Auxiliary-K. The third comparison method combines the estimated tumor mixing proportion obtained from *π*-MCP with the auxiliary calibration quantity in a two-feature consensus clustering procedure [Monti et al., 2003]. We denote this method by *π*+Auxiliary-CC. Consensus clustering is implemented using the `ConsensusClusterPlus` package [Wilkerson and Hayes, 2010].

For the MCP-based methods, we fit the model over a decreasing sequence of regularization parameter *λ*^′^*s*. For each value of the tuning parameter, we obtain the estimated transformed signal 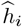. Spatial domains are then extracted from the estimated fused structure. In particular, for a given numerical tolerance *τ* ≥ 0, two neighboring spots *i* and *j* are assigned to the same domain if 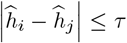. The estimated spatial domains are defined as the connected components of the graph obtained by retaining only the edges satisfying this criterion. The best solution along the regularization path is selected using the BIC criterion. For each simulated dataset, consensus clustering was performed with 1,000 resampling iterations.

We consider two simulation settings that differ in how the spatially structured auxiliary signal is related to the tumor mixing proportion. In each setting, the three domains have equal size, with (*n*_1_, *n*_2_, *n*_3_) = (50, 50, 50), yielding a total number of spots *S* = *n*_1_ + *n*_2_ + *n*_3_ = 150. We consider two gene dimensions, *G* ∈ {100, 500}. Let *d*_*i*_ ∈ {1, 2, 3}denote the true domain label of spot *i*. The auxiliary signal is generated as

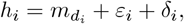

where 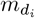 is a domain-specific mean, *ε*_*i*_ ∼ *N* (0, 0.015^2^) is independent noise, and δ = (*δ*_1_, …, *δ*_*S*_)^T^ is a smooth Gaussian process perturbation. Specifically, δ ∼ *N* (**0**, 0.02^2^**K**), with covariance kernel 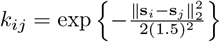, where **s**_*i*_ ∈ ℝ^2^ denotes the spatial coordinate of spot *i*.

The tumor mixing proportion is generated independently of the Gaussian process perturbation as

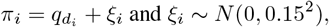

with domain-specific means (*q*_1_, *q*_2_, *q*_3_) = (0.45, 0.50, 0.55). The generated values are truncated to the interval [0.1, 0.9].

### Setting 1: multiplicative calibration. In the first setting, the domain-specific means of the auxiliary signal are

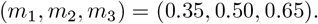

After generating *h*_*i*_ and *π*_*i*_, we define the transformation coefficients by

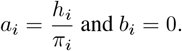

Thus, the transformed signal used in the spatial penalty is *h*_*i*_(*π*_*i*_) = *a*_*i*_*π*_*i*_.

### Setting 2: additive calibration. In the second setting, the domain-specific means of the auxiliary signal are

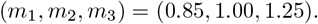

After generating *h*_*i*_ and *π*_*i*_, we define the transformation coefficients by

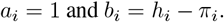

The transformed signal is therefore *h*_*i*_(*π*_*i*_) = *π*_*i*_ + *b*_*i*_.

For both transformation settings, after generating *π* and specifying the auxiliary signal, we generate the gene-specific tumor and non-tumor expression parameters. For each gene *g* = 1, …, *G*, we sample *µ*_*tg*_, *µ*_*ng*_∼ Gamma(5, 0.5), and *ϕ*_*tg*_, *ϕ*_*ng*_∼ Uniform(0.1, 10), independently across genes and components. Here *µ*_*tg*_ and *µ*_*ng*_ denote the tumor and non-tumor baseline expression levels, and *ϕ*_*tg*_ and *ϕ*_*ng*_ denote the corresponding negative binomial inverse-dispersion parameters. Conditional on these parameters, the latent tumor and non-tumor counts are generated as *T*_*ig*_∼ NB(*π*_*i*_*µ*_*tg*_, *ϕ*_*tg*_) and *N*_*ig*_ NB{ (1− *π*_*i*_)*µ*_*ng*_, *ϕ*_*ng*_}. The observed expression count is then obtained by the additive model *Y*_*ig*_ = *T*_*ig*_ + *N*_*ig*_.

Let *C*_*i*_ denote the true domain label and let 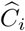 denote the estimated label for spot *i*. We evaluate domain recovery using three metrics. First, we compute the mis-clustering error, Error = Min 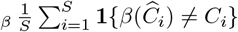, where the minimum is taken over all permutations *β* of the estimated cluster labels. Second, we compute the adjusted Rand index (ARI) [Hubert and Arabie, 1985], which measures the similarity between the estimated and true partitions after correcting for chance agreement. The ARI takes value 1 for perfect recovery, while values near 0 indicate chance-level agreement. Third, we compute the normalized variation of information (NVI) [Meil ă, 2007], an information-theoretic distance between two partitions. Smaller values of NVI indicate better agreement, with NVI equal to 0 when the two partitions are identical. For each method and simulation setting, we summarize performance across the 50 independent replications using boxplots of the clustering error rate, ARI, and NVI.

Across the boxplots in Figures 1–2, the proposed *h*(*π*)-MCP method consistently demonstrates superior spatial domain recovery compared with the three competing approaches. In all reported settings, *h*(*π*)-MCP achieves the lowest mis-clustering error, the highest ARI, and the lowest NVI. The *π*-MCP method performs poorly in most settings, especially when *G* = 500, where its misclustering error is close to 0.9 and its ARI is close to zero in Setting 1. This indicates that the spatial domain structure is not well captured by *π*_*i*_ alone. Therefore, imposing spatial smoothness directly on the model parameter may lead to inaccurate segmentation.

**Figure 1.**
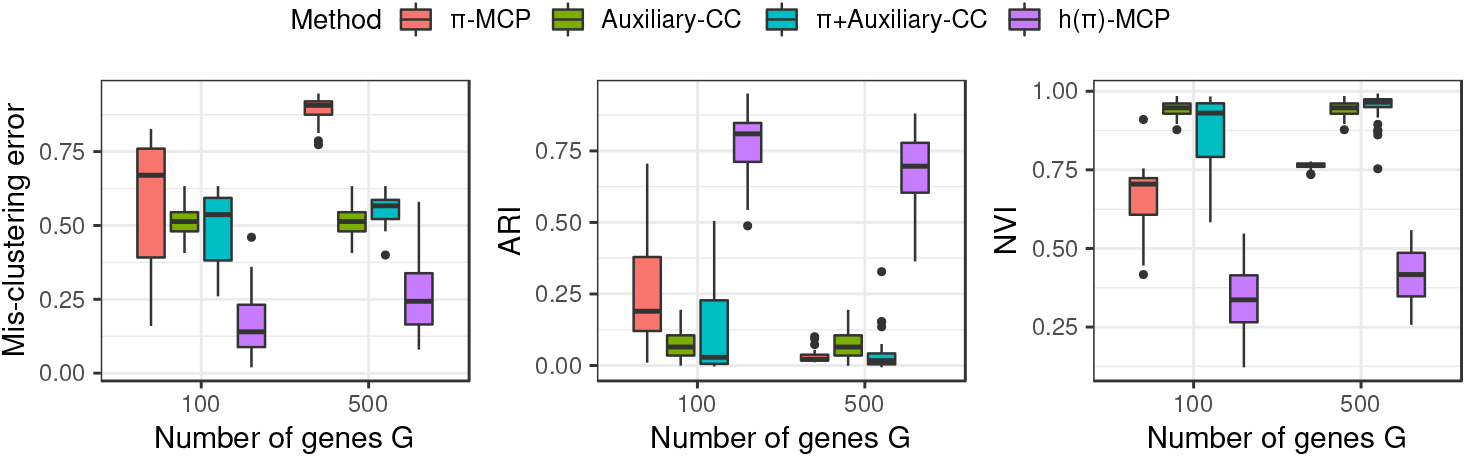
Boxplots of simulation performance for Setting 1 under the balanced domain-size scenario, with *n*_1_ = 50, *n*_2_ = 50, and *n*_3_ = 50. The three panels report the mis-clustering error, adjusted Rand index (ARI), and normalized variation of information (NVI), respectively. For each value of *G*, the boxplots compare four methods: *π*-MCP, Auxiliary-K, *π*+Auxiliary-CC, and *h*(*π*)-MCP.

**Figure 2.**
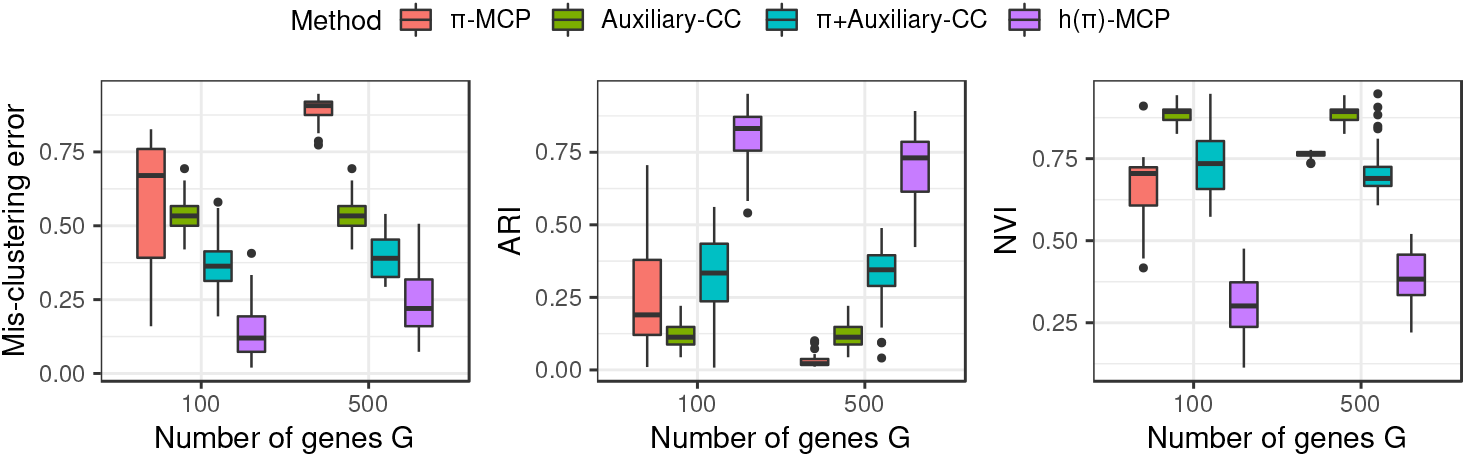
Boxplots of simulation performance for Setting 2 under the balanced domain-size scenario, with *n*_1_ = 50, *n*_2_ = 50, and *n*_3_ = 50. The three panels report the mis-clustering error, adjusted Rand index (ARI), and normalized variation of information (NVI), respectively. For each value of *G*, the boxplots compare four methods: *π*-MCP, Auxiliary-K, *π*+Auxiliary-CC, and *h*(*π*)-MCP.

Auxiliary-K, which uses only the auxiliary calibration quantity, also fails to recover the true domains reliably. Although its mis-clustering error is moderate in some imbalanced scenarios, its ARI remains low and its NVI remains high, indicating poor agreement with the true partition. This suggests that the auxiliary information alone is not sufficient for accurate domain detection. In addition, since Auxiliary-K does not use the gene expression counts, its input features remain unchanged as *G* increases. As a result, its performance remains essentially the same across different gene dimensions.

The *π*+Auxiliary-CC method improves over Auxiliary-K in several settings, particularly in Setting 2, where combining the estimated mixing proportion with the auxiliary quantity leads to a noticeable improvement in ARI and NVI. However, this two-feature consensus clustering approach remains substantially less accurate than *h*(*π*)-MCP. This comparison suggests that auxiliary information is more effective when incorporated directly into the spatially penalized likelihood, rather than used only in a downstream clustering step. The corresponding numerical summaries, including the averages and standard errors for all four simulation settings, are provided in the **Appendix**.

Figure 3 presents a spatial comparison of the clustering results for simulation Setting 1. The results indicate that incorporating auxiliary information together with *π* substantially improves clustering performance relative to using *π* alone.

**Figure 3.**
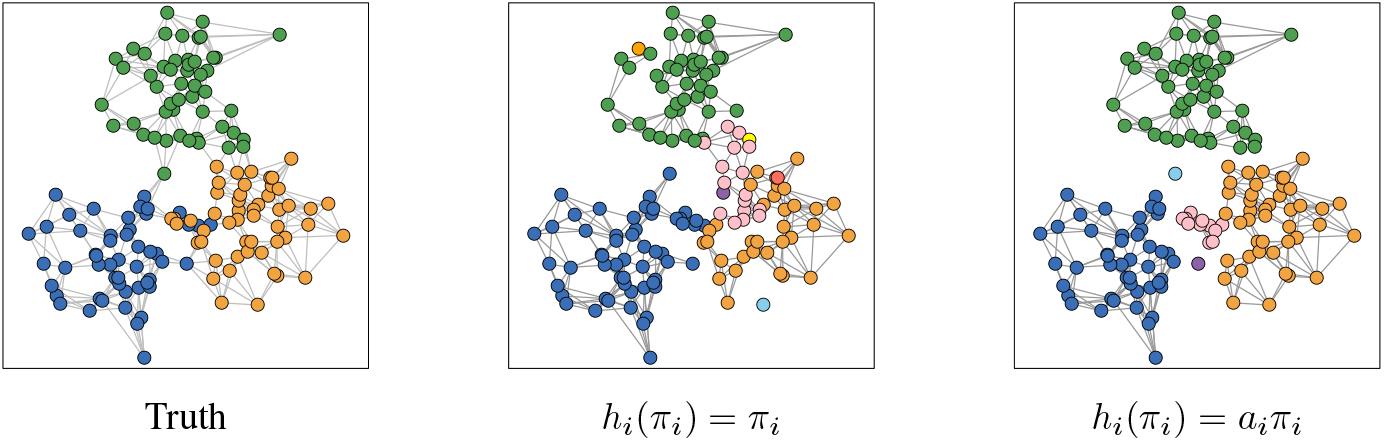
Spatial comparison of clustering results for simulation Setting 1 under the balanced domain-size scenario. The left panel shows the true domain labels, the middle panel shows clustering based only on the tumor transcript proportion *π*, and the right panel shows clustering based jointly on *π* and auxiliary information.

We further considered imbalanced domain-size scenarios under both simulation settings. The corresponding tables and plots are provided in the **Appendix**.

## 5 Discussion

A key advantage of the ATLAS framework is that auxiliary information is incorporated directly into the smoothness criterion rather than dealt with post hoc to independently estimated parameters. Our simulations demonstrate this principled integration produces sharper spatial domain recovery than two-stage strategies that first smooth and then adjust. This design is well-suited to modern biomedical studies, where spatial transcriptomics assays are routinely co-registered with complementary modalities — histology images, proteomics, chromatin accessibility, or clinical annotations — each providing a distinct but correlated view of tissue architecture. The ATLAS penalty offers the potential for a unified mechanism for any such auxiliary signal to inform smoothing without requiring it to enter the likelihood.

Although our case study focuses on spatial transcriptomics deconvolution, the ATLAS framework extends naturally to any setting where auxiliary covariates carry spatial structure. Several directions build on this foundation. Of course, local models could be adapted to different likelihood functions with minimal alteration to the ADMM algorithm used in our case study, but we see the most promising future directions, however, having to do with the design of the *h*_*i*_ transformations.

When multiple auxiliary sources are available, a natural extension of *h*_*i*_ incorporates a scalar summary of a vector of location-specific covariates ***ψ***_*i*_ = (*ψ*_*i*1_, …, *ψ*_*iK*_) — for instance, a linear combination *h*_*i*_(*π*_*i*_) = *π*_*i*_ + **c**^T^***ψ***_*i*_. In spatial transcriptomics, this would enable simultaneous fusion of histology scores, cell-type reference profiles, and imaging features into a single penalty.

Beyond multiple inputs, one may relax the affine structure on *h*_*i*_ entirely. If *h*_*i*_ is invertible and continuously differentiable with invertible Jacobian, then by the inverse function theorem *h*_*i*_ ^−1^ is differentiable. This opens the door to solving the following optimization problem using splitting techniques like the proposed ADMM algorithm.

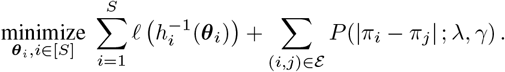

More generally though, if *h*_*i*_ is not invertible, it is possible to approximate with high accuracy the proximal operator of the composition penalty using Monte Carlo techniques Osher et al. [2023], Tibshirani et al. [2025]. These approximations can be substituted into splitting methods like our ADMM implementation without compromising global convergence guarantees [Di et al., 2026]. These extensions suggest ATLAS can be deployed broadly, wherever auxiliary spatial structure is available and a closed-form proximal operator is unavailable.

Finally, the above directions assume *h*_*i*_ has a pre-specified structure, but we could consider fitting a shallow neural network to learn in a data-driven manner how to fold auxiliary information into the smoothing criterion. While important details need be determined, the flexibility of *h*_*i*_, whether pre-specified or learned, positions ATLAS as a general scaffold for principled multimodal integration in spatially structured estimation problems.

## S1 Appendix

### S1.1 Real-Data Motivation

Figure S1 compares the spatial clustering patterns obtained from the tumor transcript proportion *π*, the tumor cell purity *ρ*, and their difference *π*−*ρ* using lung squamous cell carcinoma (LUSC) sample [Montierth et al., 2025].

**Figure S1.**
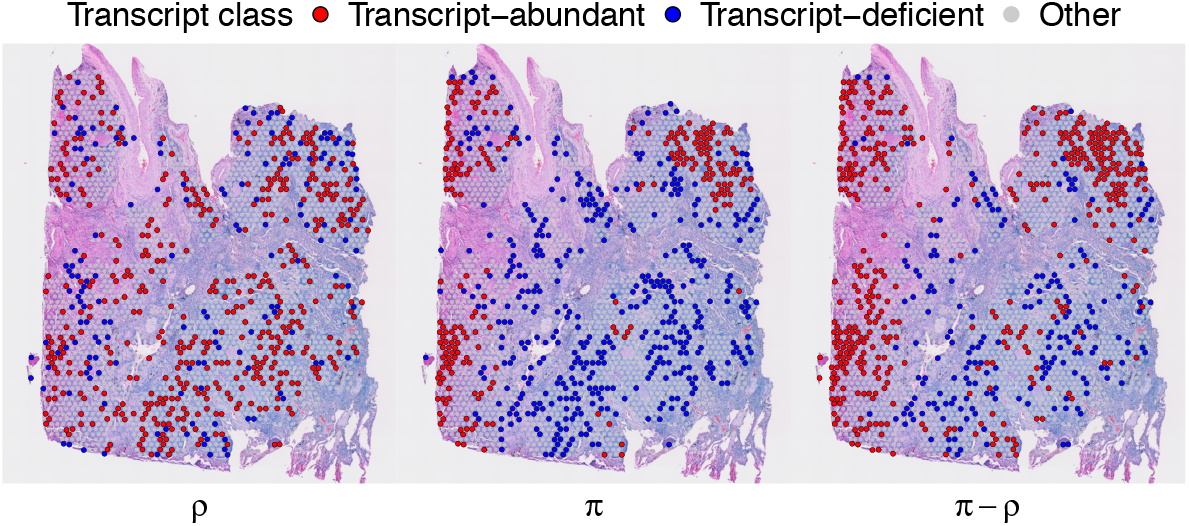
Spatial comparison of clustering patterns in a lung squamous cell carcinoma (LUSC) sample. The left panel shows k-mean clustering based on the tumor cell purity *ρ*, the middle panel shows clustering based on the tumor transcript proportion *π*, and the right panel shows clustering based on their difference *π* − *ρ*.

### S1.2 Algorithmic Details

**Update of *π*** We provide additional details for the Newton update used in the ***π***-step of the ADMM algorithm. At ADMM iteration *k*, the ***π***-subproblem decomposes into *S* independent scalar optimization problems. For spot *i*, the objective is

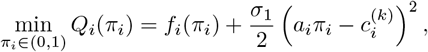

Where 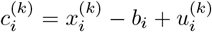. To enforce the constraint *π*_*i*_ ∈ (0, 1), we reparameterize the scalar problem using the logit transformation 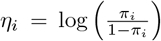, and 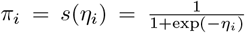. The optimization is then performed over the unconstrained variable *η*_*i*_.

Let 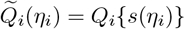. The Newton direction is computed as 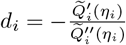. By the chain rule,

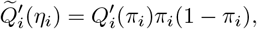

and

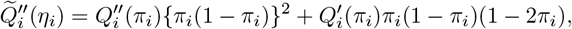

Where

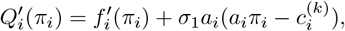

and

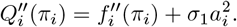

Here 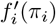 and 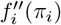 are obtained by differentiating the spot-specific convolution log-likelihood.

We choose a step size *α* ∈(0, 1] by Armijo backtracking line search. That is, starting from *α* = 1, we repeatedly shrink *α* until

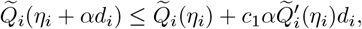

where *c*_1_ ∈ (0, 1) is the Armijo parameter. The inner iteration is updated by

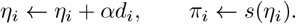

#### Algorithm 1

ADMM algorithm for transformed MCP spatial smoothing

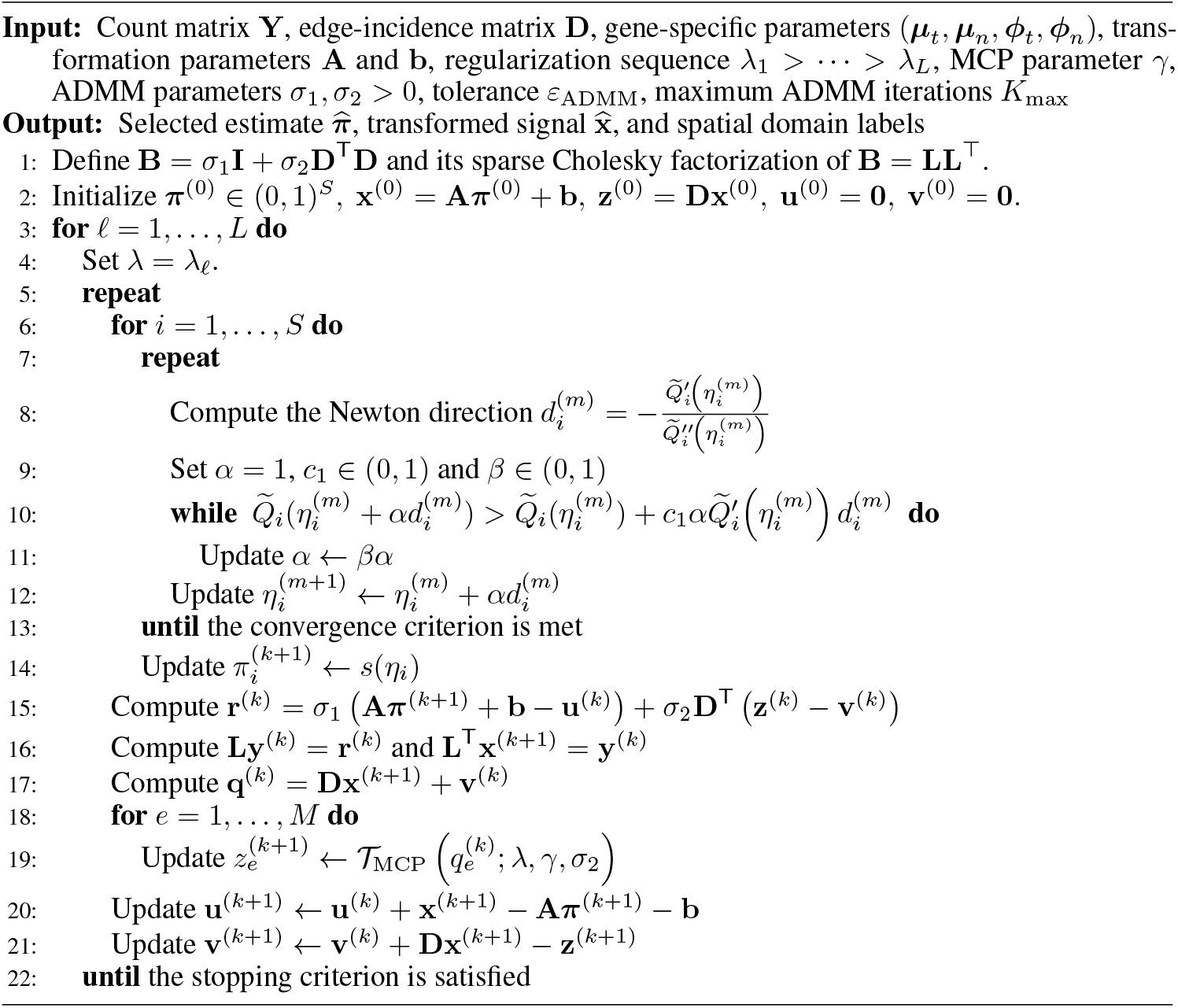

**Update of x** We provide additional details on the sparse linear algebra used in the **x**-update. At ADMM iteration *k*, the update of **x** requires solving the linear system

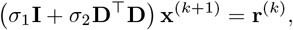

Where

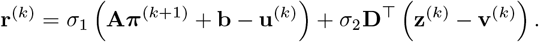

Let **B** = *σ*_1_**I** + *σ*_2_**D**^T^**D**. Since **D**^T^**D** is the graph Laplacian associated with the spatial adjacency graph, it is positive semidefinite. Therefore, when *σ*_1_ *>* 0, the shifted matrix **B** is symmetric positive definite, and the linear system has a unique solution.

We solve this system using a sparse Cholesky factorization. Specifically, we compute

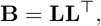

where **L** is a sparse lower triangular matrix. The solution is then obtained by two triangular solves:

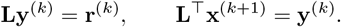

For fixed *σ*_1_, *σ*_2_, and spatial graph, the matrix **B** does not depend on the ADMM iteration index *k* or on the regularization parameter *λ*. Thus, its sparse Cholesky factorization can be computed once and reused throughout the ADMM iterations. When the model is fit over a regularization path, the same factorization can also be reused across all values of *λ*, so that only the right-hand side **r**^(*k*)^ needs to be updated at each iteration.

The algorithm is summarized in Algorithm 1.

**Figure S2.**
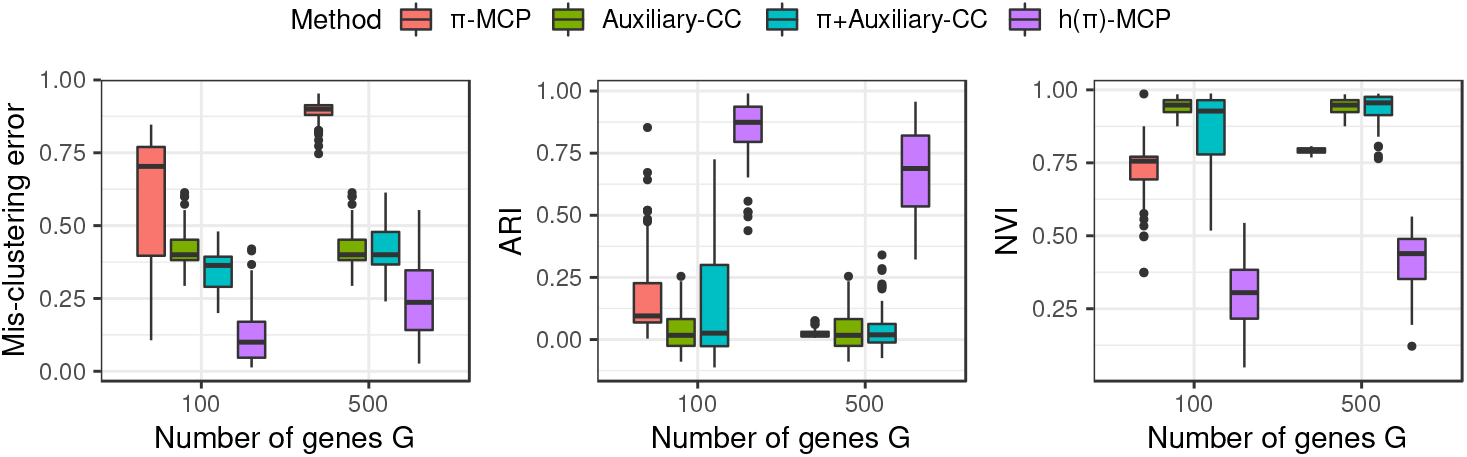
Boxplots of simulation performance for Setting 1 under the imbalanced domain-size scenario, with *n*_1_ = 30, *n*_2_ = 30, and *n*_3_ = 90. The three panels report the mis-clustering error, adjusted Rand index (ARI), and normalized variation of information (NVI), respectively. For each value of *G*, the boxplots compare four methods: *π*-MCP, Auxiliary-K, *π*+Auxiliary-CC, and *h*(*π*)- MCP. Each boxplot is based on 50 independent simulation runs.

**Figure S3.**
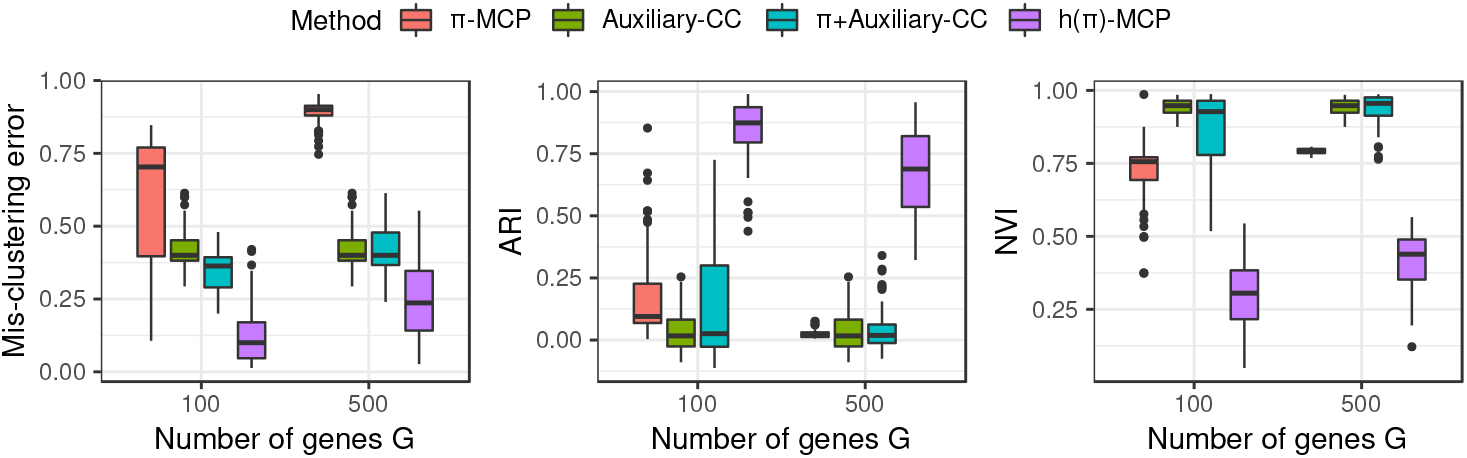
Boxplots of simulation performance for Setting 2 under the imbalanced domain-size scenario, with *n*_1_ = 30, *n*_2_ = 30, and *n*_3_ = 90. The three panels report the mis-clustering error, adjusted Rand index (ARI), and normalized variation of information (NVI), respectively. For each value of *G*, the boxplots compare four methods: *π*-MCP, Auxiliary-K, *π*+Auxiliary-CC, and *h*(*π*)- MCP. Each boxplot is based on 50 independent simulation runs.

### S1.3 Additional Simulation Results

Tables S2–S4 summarize the numerical results for the two simulation settings under the imbalanced domain-size scenario, where *n*_1_ = 30, *n*_2_ = 30, and *n*_3_ = 90. Across all cases, the proposed *h*(*π*)-MCP method achieves the best overall performance, attaining the lowest mis-clustering error, the highest ARI, and the lowest NVI for both *G* = 100 and *G* = 500. This pattern is observed under both balanced and imbalanced domain-size scenarios, suggesting that the proposed method remains effective even when one spatial domain is substantially larger than the others.

**Table S1.** Simulation results for Setting 1 under the balanced domain-size scenario, with *n*_1_ = 50, *n*_2_ = 50, and *n*_3_ = 50. The table reports the mis-clustering error, adjusted Rand index (ARI), and normalized variation of information (NVI) for *π*-MCP, Auxiliary-K, and *π*+Auxiliary-CC and *h*(*π*)-MCP. The numbers are the average quantities over 50 independent runs and the standard errors are presented in the parentheses.

| | | π-MCP | Auxiliary-K | $\pi$+Auxiliary-CC | $h(\pi)$-MCP |
| --- | --- | --- | --- | --- | --- |
| G=100 | Error | 0.581<sub>(0.031)</sub> | 0.517<sub>(0.007)</sub> | 0.499<sub>(0.016)</sub> | 0.162<sub>(0.014)</sub> |
|  | ARI | 0.275<sub>(0.029)</sub> | 0.072<sub>(0.007)</sub> | 0.132<sub>(0.024)</sub> | 0.782<sub>(0.016)</sub> |
|  | NVI | 0.664<sub>(0.014)</sub> | 0.944<sub>(0.003)</sub> | 0.872<sub>(0.017)</sub> | 0.337<sub>(0.015)</sub> |
| G=500 | Error | 0.897<sub>(0.005)</sub> | 0.517<sub>(0.007)</sub> | 0.557<sub>(0.006)</sub> | 0.253<sub>(0.016)</sub> |
|  | ARI | 0.030<sub>(0.003)</sub> | 0.072<sub>(0.007)</sub> | 0.034<sub>(0.008)</sub> | 0.684<sub>(0.017)</sub> |
|  | NVI | 0.763<sub>(0.001)</sub> | 0.944<sub>(0.003)</sub> | 0.953<sub>(0.006)</sub> | 0.413<sub>(0.011)</sub> |

**Table S2.** Simulation results for Setting 1 under the imbalanced domain-size scenario, with *n*_1_ = 30, *n*_2_ = 30, and *n*_3_ = 90. The table reports the mis-clustering error, adjusted Rand index (ARI), and normalized variation of information (NVI) for *π*-MCP, Auxiliary-K, and *π*+Auxiliary-CC and *h*(*π*)-MCP. The numbers are the average quantities over 50 independent runs and the standard errors are presented in the parentheses.

| | | π-MCP | Auxiliary-K | $\pi$+Auxiliary-CC | $h(\pi)$-MCP |
| --- | --- | --- | --- | --- | --- |
| G=100 | Error | 0.597<sub>(0.032)</sub> | 0.425<sub>(0.011)</sub> | 0.342<sub>(0.011)</sub> | 0.124<sub>(0.004)</sub> |
|  | ARI | 0.195<sub>(0.029)</sub> | 0.031<sub>(0.012)</sub> | 0.130<sub>(0.031)</sub> | 0.834<sub>(0.020)</sub> |
|  | NVI | 0.716<sub>(0.016)</sub> | 0.943<sub>(0.004)</sub> | 0.865<sub>(0.019)</sub> | 0.306<sub>(0.016)</sub> |
| G=500 | Error | 0.889<sub>(0.006)</sub> | 0.425<sub>(0.011)</sub> | 0.419<sub>(0.011)</sub> | 0.245<sub>(0.017)</sub> |
|  | ARI | 0.024<sub>(0.002)</sub> | 0.031<sub>(0.012)</sub> | 0.048<sub>(0.013)</sub> | 0.680<sub>(0.025)</sub> |
|  | NVI | 0.792<sub>(0.001)</sub> | 0.943<sub>(0.004)</sub> | 0.938<sub>(0.008)</sub> | 0.415<sub>(0.015)</sub> |

**Table S3.** Simulation results for Setting 2 under the balanced domain-size scenario, with *n*_1_ = 50, *n*_2_ = 50, and *n*_3_ = 50. The table reports the mis-clustering error, adjusted Rand index (ARI), and normalized variation of information (NVI) for *π*-MCP, Auxiliary-K, and *π*+Auxiliary-CC and *h*(*π*)-MCP. The numbers are the average quantities over 50 independent runs and the standard errors are presented in the parentheses.

| | | π-MCP | Auxiliary-K | $\pi$+Auxiliary-CC | $h(\pi)$-MCP |
| --- | --- | --- | --- | --- | --- |
| G=100 | Error | 0.581<sub>(0.031)</sub> | 0.427<sub>(0.007)</sub> | 0.369<sub>(0.012)</sub> | 0.140<sub>(0.013)</sub> |
|  | ARI | 0.275<sub>(0.029)</sub> | 0.171<sub>(0.007)</sub> | 0.320<sub>(0.020)</sub> | 0.811<sub>(0.014)</sub> |
|  | NVI | 0.664<sub>(0.014)</sub> | 0.882<sub>(0.004)</sub> | 0.738<sub>(0.013)</sub> | 0.303<sub>(0.014)</sub> |
| G=500 | Error | 0.897<sub>(0.005)</sub> | 0.535<sub>(0.008)</sub> | 0.394<sub>(0.010)</sub> | 0.249<sub>(0.016)</sub> |
|  | ARI | 0.030<sub>(0.003)</sub> | 0.119<sub>(0.006)</sub> | 0.324<sub>(0.013)</sub> | 0.695<sub>(0.017)</sub> |
|  | NVI | 0.763<sub>(0.001)</sub> | 0.886<sub>(0.003)</sub> | 0.709<sub>(0.010)</sub> | 0.390<sub>(0.011)</sub> |

**Table S4.** Simulation results for Setting 2 under the imbalanced domain-size scenario, with *n*_1_ = 30, *n*_2_ = 30, and *n*_3_ = 90. The table reports the mis-clustering error, adjusted Rand index (ARI), and normalized variation of information (NVI) for *π*-MCP, Auxiliary-K, and *π*+Auxiliary-CC and *h*(*π*)-MCP. The numbers are the average quantities over 50 independent runs and the standard errors are presented in the parentheses.

| | | π-MCP | Auxiliary-K | $\pi$+Auxiliary-CC | $h(\pi)$-MCP |
| --- | --- | --- | --- | --- | --- |
| G=100 | Error | 0.597<sub>(0.032)</sub> | 0.541<sub>(0.008)</sub> | 0.285<sub>(0.011)</sub> | 0.145<sub>(0.015)</sub> |
|  | ARI | 0.195<sub>(0.029)</sub> | 0.134<sub>(0.007)</sub> | 0.430<sub>(0.024)</sub> | 0.800<sub>(0.024)</sub> |
|  | NVI | 0.716<sub>(0.016)</sub> | 0.878<sub>(0.004)</sub> | 0.699<sub>(0.016)</sub> | 0.307<sub>(0.018)</sub> |
| G=500 | Error | 0.889<sub>(0.006)</sub> | 0.541<sub>(0.008)</sub> | 0.361<sub>(0.014)</sub> | 0.230<sub>(0.018)</sub> |
|  | ARI | 0.024<sub>(0.002)</sub> | 0.134<sub>(0.007)</sub> | 0.509<sub>(0.023)</sub> | 0.697<sub>(0.025)</sub> |
|  | NVI | 0.792<sub>(0.001)</sub> | 0.878<sub>(0.004)</sub> | 0.673<sub>(0.014)</sub> | 0.398<sub>(0.016)</sub> |

The boxplots in Figures S2–S3 further illustrate that the proposed *h*(*π*)-MCP method consistently provides superior spatial domain recovery compared with the three competing approaches.

